# Quantitative single-cell interactomes in normal and virus-infected mouse lungs

**DOI:** 10.1101/2020.02.05.936054

**Authors:** Margo P Cain, Belinda J Hernandez, Jichao Chen

## Abstract

Mammalian organs consist of diverse, intermixed cell types that signal to each other via ligand-receptor interactions – an interactome – to ensure development, homeostasis, and injury-repair. Dissecting such intercellular interactions is facilitated by rapidly growing single-cell RNA-seq (scRNA-seq) data; however, existing computational methods are often not sufficiently quantitative nor readily adaptable by bench scientists without advanced programming skills. Here we describe a quantitative intuitive algorithm, coupled with an optimized experimental protocol, to construct and compare interactomes in control and Sendai virus-infected mouse lungs. A minimum of 90 cells per cell type compensates for the known gene dropout issue in scRNA-seq and achieves comparable sensitivity to bulk RNA-seq. Cell lineage normalization after cell sorting allows cost-efficient representation of cell types of interest. A numeric representation of ligand-receptor interactions identifies, as outliers, known and potentially new interactions as well as changes upon viral infection. Our experimental and computational approaches can be generalized to other organs and human samples.

**Summary statement:** An intuitive method to construct quantitative ligand-receptor interactomes using single-cell RNA-seq data and its application to normal and Sendai virus-infected mouse lungs.

## INTRODUCTION

In multicellular mammalian organs, specialized cell types, such as beta cells in the pancreas and cardiomyocytes in the heart, perform organ-specific functions in coordination with generic cell types, such as the omnipresent endothelial and immune cells. Such coordination occurs at the microscopic level such that many organs can be viewed as basic units repeated millions of times, as exemplified by alveoli in the lung, glomeruli in the kidney, and hair follicles in the skin. Beside physiology, these multi-cell type units are also integrated niches underlying homeostatic maintenance and injury-repair (Chang-Panesso and Humphreys, 2017; Tata and Rajagopal, 2017; Varga and Greten, 2017). On the molecular level, cell-cell communication is chiefly mediated by secreted ligands and membrane receptors, and is made possible by continuous patrolling of immune cells and more permanent contacts via interdigitating cellular processes embedded in extracellular matrices. This can be readily appreciated in the lung, where each alveolus is mostly air bordered by several intermixed cells of the epithelial, endothelial, mesenchymal, and immune cell lineages such that most, if not all, cell types are within a signaling distance of each other (Cohen et al., 2018; Raredon et al., 2019; Yuan et al., 2018)

Delineating genome-wide ligand-receptor interactions between all pairwise combinations of constituent cell types in a given organ, hereinafter named interactomes, becomes feasible with the advancement of single-cell RNA-seq (scRNA-seq) technology, where individual cell types can be profiled upon computational, instead of physical, purification (Han et al., 2018; Tabula Muris et al., 2018)). Several such single-cell interactomes have been constructed to characterize organs and cell culture systems at baseline and upon perturbation (Camp et al., 2017; Cohen et al., 2018; Kumar et al., 2018; Raredon et al., 2019; Skelly et al., 2018; Vento-Tormo et al., 2018; Vieira Braga et al., 2019). Most studies apply a threshold to categorize ligands and receptors as present or absent and count these binary outcomes as a measure of interaction strength between cell types – a missed opportunity given the numerical expression values available from scRNA-seq. Furthermore, systematic comparison of scRNA-seq and bulk RNA-seq is often not incorporated into experimental design to allay the known gene dropout concern in scRNA-seq. Finally, current algorithms and outputs are not readily adopted by bench scientists without advanced computational skills.

In this methodological study using the mouse lung as a model system, we have evaluated the sensitivity and cellular resolution of scRNA-seq in comparison with bulk RNA-seq, accordingly optimized a cell sorting protocol to consistently capture all major lung cell types, and generated numeric interactomes in normal and diseased lungs where significant interactions are intuitively represented as outliers. Developed by bench scientists, our combined experimental and computational methods can be generalized to other biological systems with minimal adjustment.

## RESULTS

### ScRNA-seq has cell type resolution with sensitivity comparable to bulk RNA-seq

One challenge to construct an interactome using scRNA-seq is the so-called gene dropout issue where only a few thousand genes are detected in a given cell due to technical inefficiency, as compared to the 20,000-30,000 genes expected and obtained by bulk RNA-seq of typical mammalian cells (Hicks et al., 2018; Kharchenko et al., 2014). However, it is important to recognize that bulk RNA-seq does not detect all genes in every cell of the sample and the apparent high coverage is the result of summing over tens of thousands of cells. We thus hypothesized that, by combining cells of a cell type readily identifiable in scRNA-seq, we could achieve comparable sensitivity to bulk RNA-seq of the same purified cell type. To test this, we used as a standard our published bulk RNA-seq data of FACS-purified alveolar type 1 (AT1) and alveolar type 2 (AT2) cells (Little et al., 2019) and evaluated scRNA-seq gene dropouts as a function of expression level (Fig. 1A and S1A). We found that the dropout rate, defined as the percentage of genes detected by bulk RNA-seq but not scRNA-seq, decreased with increasing expression levels, as measured by fragments per kilobase of transcript per million mapped reads (FPKM) from bulk RNA-seq. For example, only 4% of genes at 10 FPKM were missed in scRNA-seq of 345 AT1 cells and this percentage was even lower (0.6%) for AT2 cells where 1,252 cells were sequenced. Technical or biological noise likely dominated for low FPKM genes as 61 and 54 genes were detected by scRNA-seq but not bulk RNA-seq for AT1 and AT2 cells, respectively. The consistency between scRNA-seq and bulk RNA-seq was further supported by the good agreement between the levels of gene expression measured by both methods (Fig. 1B, Table S1).

**Figure 1.**
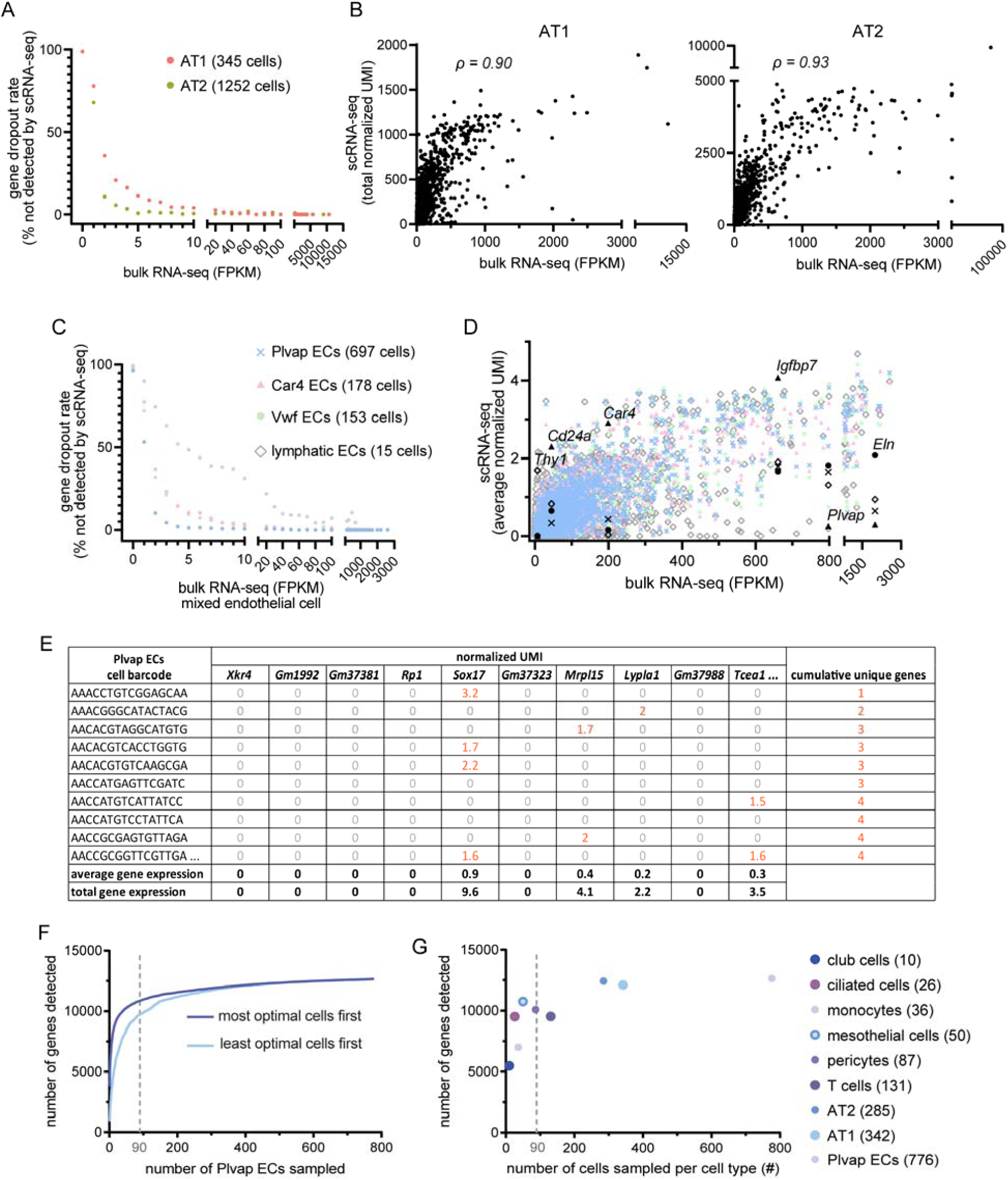
Cell type-level scRNA-seq data compares favorably to bulk RNA-seq. (**A**) scRNA-seq gene dropout rates of AT1 and AT2 cells, with the number of cells sequenced in parenthesis, as a function of bulk RNA-seq expression (FPKM) of the corresponding FACS-purified cell types. The dropout rate is the percentage of genes not detected by scRNA-seq for a given range of bulk RNA-seq expression values. ScRNA-seq data are from postnatal day (P) 7 (GSE144769); bulk RNA-seq data are from P5 for AT1 cells and P8 for AT2 cells, as published (Little et al., 2019). (**B**) Consistency between AT1 cell (left) and AT2 cell (right) gene expression from scRNA-seq and bulk RNA-seq gene expression of the corresponding cell type. The scRNA-seq equivalent of the FPKM value in bulk RNA-seq, which sums the expression in all cells of the sample, is considered the sum of normalized unique molecular identifiers (UMIs normalized by the sequencing depth of each cell) over all cells in the corresponding cell type. ρ, Spearman’s rank correlation coefficient. (**C**) scRNA-seq gene dropout rates of endothelial cell (EC) subpopulations at P7, with the number of cells sequenced in parenthesis, as a function of bulk RNA-seq expression (FPKM) of FACS-purified total ECs at P7, as published (Vila Ellis et al., 2019). (**D**) ScRNA-seq average gene expression for the 4 EC subpopulations, color/symbol-coded as in (**C**), as a function of bulk RNA-seq gene expression of total ECs. Over a wide range of FPKM values, subpopulation-specific genes are readily identifiable in scRNA-seq. Examples are marked with the same subpopulation symbols but in black. *Thy1* is enriched in lymphatic ECs; *Cd24a, Car4*, and *Igfbp7* are enriched in Car4 ECs; *Plvap* is depleted in Car4 ECs; *Eln* is enriched in Plvap ECs. One outlying, non-differential gene (*Malat1*; Table S1) is removed to decompress the plot. (**E**) Table showing the first 10 genes for the first 10 cells to illustrate how average and total gene expression values are calculated. Nonzero values are highlighted in red and tallied to obtain the number of cumulative unique genes. (**F**) The number of genes detected, as tallied in (**E**), plateaus as a function of the number of cells sequenced for a given cell type (Plvap ECs of adult lungs). Cells are sorted in ascending (light blue) or descending (dark blue) order in (**E**) by the number of genes detected in the cell. The dashed gray line marks 90 cells. (**G**) The number of genes detected for indicated cell types with the number of cells sequenced in parenthesis. The dashed gray line marks 90 cells. (**E**-**G**) use scRNA-seq data from the adult lung (GSE144678)

After establishing the comparable performance of scRNA-seq and bulk RNA-seq in measuring the same cell populations, we examined the added benefit of scRNA-seq in dissecting cell type heterogeneity without prior knowledge and availability of surface markers or genetic drivers that were necessary for physical purification in bulk RNA-seq. For this, we used lung endothelial cells (ECs) that were bulk purified with a pan-EC marker ICAM2 but contained 4 subpopulations in scRNA-seq: Plvap ECs, Car4 ECs, Vwf ECs, and lymphatic ECs (Vila Ellis et al., 2019) (Fig. S1B). As before, dropout rates were low except for lymphatic ECs, probably because only 15 cells were sequenced (Fig. 1C). However, the apparent higher sensitivity of bulk RNA-seq was likely driven by expression of those dropout genes in non-lymphatic ECs and not because bulk RNA-seq detected mRNAs derived from lymphatic ECs. Nevertheless, cell type heterogeneity captured by scRNA-seq was demonstrated by comparing population-specific gene expression as a function of expression in bulk RNA-seq (Fig. 1D, Table S1). Again as before, the majority of the genes showed good agreement between scRNA-seq and bulk RNA-seq. As we hypothesized, cell type-specific genes over a range of expression levels, such as *Cd24a, Car4*, and *Igfbp7* for Car4 ECs, showed the expected enrichment; the converse was also true for genes depleted in Car4 ECs, such as *Plvap*. Notably, *Thy1*, a known lymphatic EC marker (Jurisic et al., 2010), was negligible in bulk RNA-seq but readily detected and distinguishable in the lymphatic ECs of only 15 cells (Fig. 1D). Such cell type stratification was, by design, lost in bulk RNA-seq.

The varying performance of scRNA-seq seemingly as a function of cell number prompted a systematic evaluation: we computationally sampled the Plvap EC population (Fig. 1E) and found that the number of detected genes rapidly approached the technical limit such that ∼90 cells were necessary to detect 10,000 genes (Fig. 1F). The same 90-cell cutoff held true when comparing cell types of different cell numbers (Fig. 1G) and therefore was used to guide the optimization of our scRNA-seq wet-lab protocol as described below.

Overall, our analyses showed that scRNA-seq performs as well as bulk RNA-seq in detecting and quantifying genes when computationally combining enough cells for a given cell type, and outperforms bulk RNA-seq in identifying cell type heterogeneity and associated marker genes.

### An optimized normalized lung scRNA-seq protocol

The theoretical minimal number of cells, as determined above, could be difficult to obtain in practice, due to highly skewed proportions of the dozens of cell types in a typical mammalian organ. For example, published lung scRNA-seq datasets had limited representation of endothelial and mesenchymal cells and significant variations in cell proportions across experiments (Fig. 2A). We reasoned that consistent and sufficient sampling of major lung cell types could be achieved by first purifying and then sequencing equal proportion of the 4 cell lineages – epithelial, endothelial, immune, and mesenchymal lineages, which were generally considered non-interconvertible. As a benchmark, we determined via immunostaining the in vivo average proportions of the 4 listed cell lineages as 26%, 38%, 17%, and 19% – a skewed and variable distribution that warranted consideration in experimental design (Fig. 2B). We then identified 3 cell surface markers that robustly distinguished the 4 lineages in FACS and, in comparison with our immunostaining results, introduced biases presumably due to varying efficiency in dissociating cells of different lineages (Fig. 2C). To reduce the cost of scRNA-seq, we remixed and sequenced equal numbers of cells from the purified 4 lineages after taking into account lineage-specific difference in cell viability (Fig. 2C).

**Figure 2:**
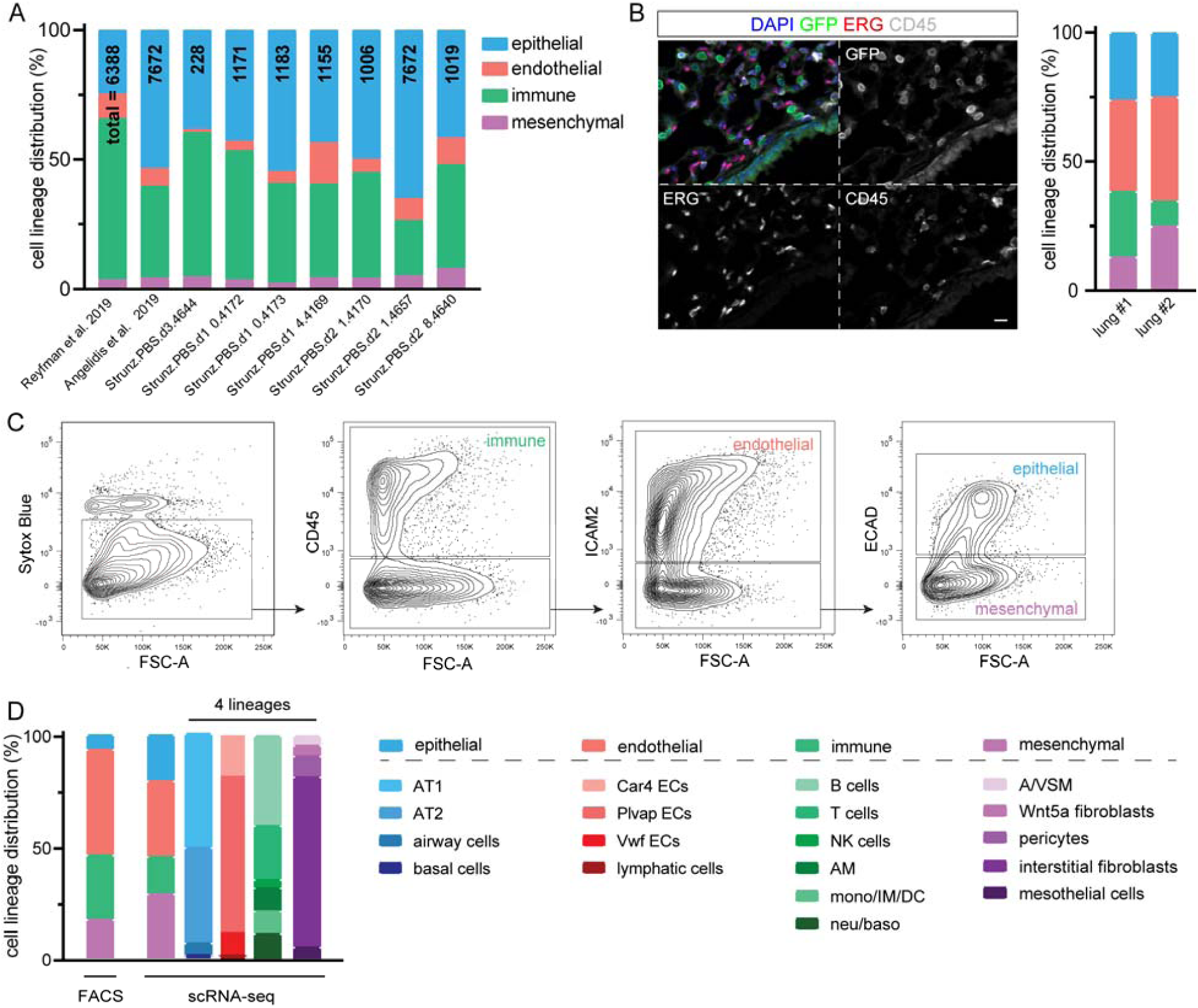
Optimized sample preparation protocol for scRNA-seq captures major lung cell types of the epithelial, endothelial, immune, and mesenchymal lineages. (**A**) Distribution of the 4 color-coded lineages quantified from published whole lung scRNA-seq datasets (Angelidis et al., 2019; Reyfman et al., 2019; Strunz et al., 2019). (**B**) Confocal images of immunostained adult *Rosa*^*Sun1GFP/+*^; *Shh*^*Cre/+*^ lungs where epithelial cell nuclei are genetically marked by nuclear envelope-targeted GFP (Mo et al., 2015), whereas ERG and CD45 mark endothelial and immune cells, respectively, and triple negative nuclei (DAPI) are considered mesenchymal. We used the GFP reporter instead of NKX2-1 because both NKX2-1 and ERG are rabbit antibodies. Percentages are from 2 lungs with 3 images each containing thousands of cells. Scale: 10 um. (**C**) An all-inclusive FACS gating strategy to separate all live cells (Sytox Blue negative) into the 4 lung cell lineages. (**D**) Skewed distributions of the 4 color-coded lung cell lineages from FACS are compensated by remixing them in equal proportions, adjusted for lineage-specific cell viability, for scRNA-seq. 3,245 cells were sequenced. Distributions of the constituent cell types in each lineage can be obtained from scRNA-seq. airway cells: ciliated and club cells; NK cells: natural killer cells; AM: alveolar macrophages; mono: monocytes; IM: interstitial macrophages; DC: dendritic cells; neu: neutrophils; baso: basophils; A/VSM: airway/vascular smooth muscle cells.

This cell-lineage-level normalization was a cost-effective trade-off between non-selective whole lung scRNA-seq and in-depth albeit narrow-focused cell type-specific scRNA-seq. Proportions of cell lineages and individual cell types within a lineage could be retrieved by analyzing FACS and scRNA-seq data, respectively (Fig. 2D). Our method routinely captured 18 lung cell types in a sufficient number to construct the interactome.

### Numeric representation of ligand-receptor interaction

As ligand-receptor interaction was directional – consisting of ligand-expressing signaling cells and receptor-expressing receiving cells, we evaluated each cell type in our scRNA-seq for its potential as ligand-expressing cells when paired with each of all cell types, including itself in the case of autocrine interaction (Fig. 3A, Table S2). For each of these directional cell type pairs, we used a scatterplot to visualize all 2,356 ligand-receptor pairs such that a data point off both axes indicated the presence of the corresponding ligand and receptor, as exemplified by the expected *Vegfa*-*Kdr* expression in the AT1 cell-Car4 EC pair (Vila Ellis et al., 2019; Yang et al., 2016) (Fig. 3A). In these scatterplots, user-defined horizontal and vertical thresholds could be used to tally all ligand-receptor pairs present in specific cell type pairs – an approach commonly employed in the literature but at the expense of available quantitative expression values (Camp et al., 2017; Cohen et al., 2018; Skelly et al., 2018). Furthermore, the number of ligand-receptor pairs was not necessarily a valid predictor of functional interactions, and a single threshold was unlikely to suit all ligands and receptors expressed at varied levels.

**Figure 3:**
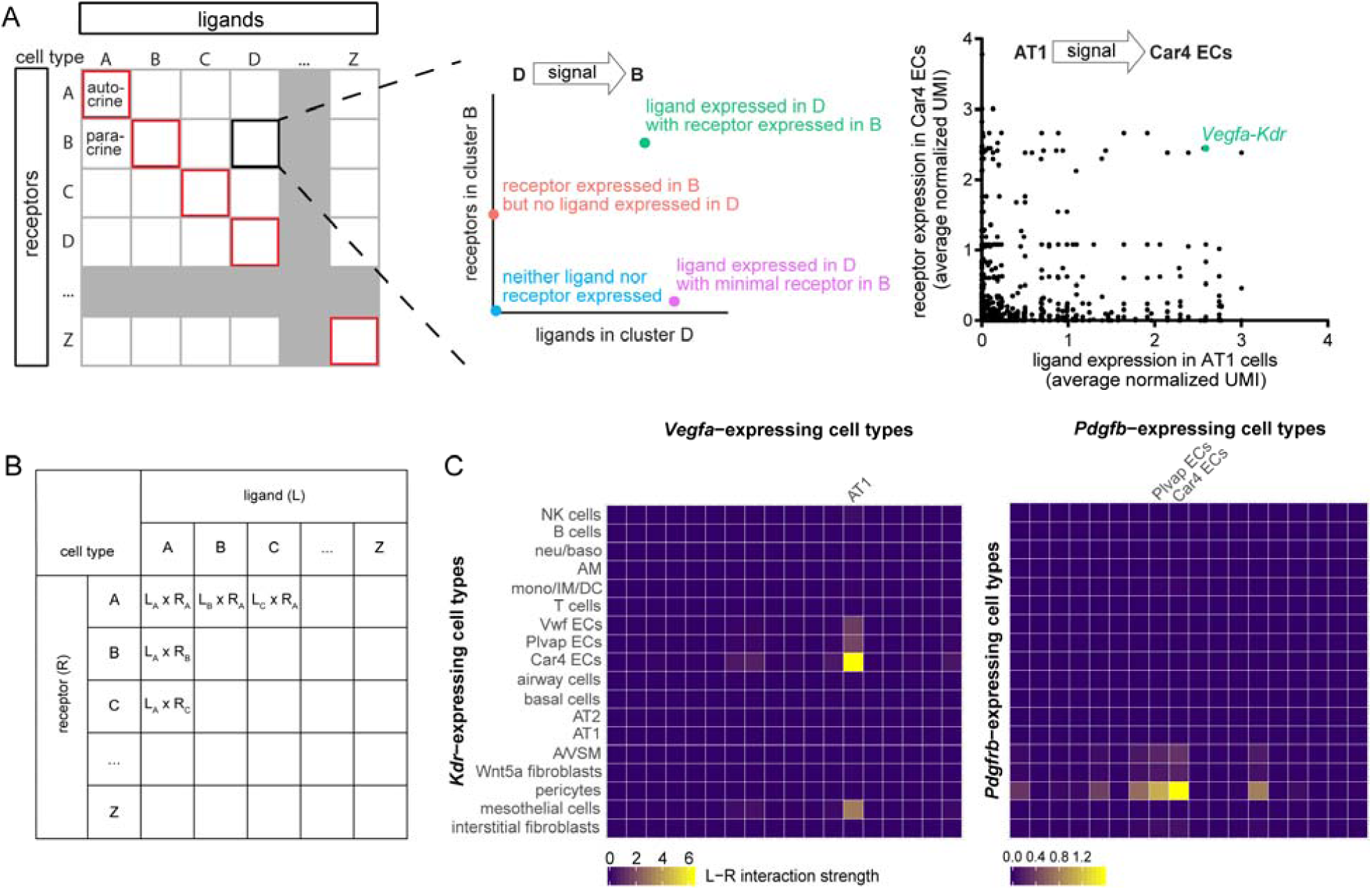
Constructing interactomes using numerical representations of ligand-receptor interaction capitalizes on quantitative information from scRNA-seq. (**A**) Schematic illustrating ligand-receptor interactions between directional cell type-pairs. Autocrine (between the same cell type albeit not necessarily the same cell) interactions are boxed in red along the diagonal. A hypothetical cell type-pair (D to B) illustrates the locations of possible ligand-receptor interactions, as demonstrated in a real example showing a known *Vegfa*-*Kdr* interaction from AT1 cells to Car4 ECs. (**B**) Table illustrating that for a given ligand-receptor pair, the interaction strength for each directional cell type-pair is defined as the product of the average ligand expression in the signaling cells and the average receptor expression in the receiving cells. (**C**) Heatmaps to visualize the hypothetical table in (**B**) for 2 real ligand-receptor interactions known to occur between AT1 cells and Car4 ECs via *Vegfa*-*Kdr* and between ECs and pericytes via *Pdgfb*-*Pdgfrb*. The cell types for the columns are not fully labeled but are in the same order as those for the rows, as in (**B**).

To overcome these limitations, we sought a numeric representation of ligand-receptor interaction, which we reasoned should positively correlate with the expression values of both the ligand and the receptor. Inspired by the principle of chemical equilibrium where the product –the ligand-receptor complex in this case, which should determine the signaling output – equals the product of the substrates divided by the equilibrium constant, we chose to represent the interaction strength by multiplying the average scRNA-seq expression values of the ligand and the receptor in the involved cell types (Fig. 3B). By assuming comparable protein translation and delivery among cell types, this numeric representation allowed us to leverage 324 cell type pairs for a given ligand-receptor pair and readily captured potential interactions as outliers. This was demonstrated using heatmaps for the known interactions between AT1 cells and Car4 ECs via *Vegfa* and *Kdr*, and between ECs and pericytes via *Pdgfb* and *Pdgfrb* (Fig. 3C). These visually outlying interactions were quantitatively defined as those outside 3 times the interquartile range (Table S3, S4).

### Identify interactions altered upon viral infection

Next, we extended our numeric interactome analysis to compare lungs at baseline and upon perturbation. We chose a Sendai virus infection model (Goldblatt et al., 2020), instead of genetic models, because infection was expected to induce global changes involving multiple cell types and their associated cell interactions, necessitating a quantitative genomic analysis as our interactome method. At 2 weeks after infection, when the virus had been largely cleared and the lung was undergoing repair (Goldblatt et al., 2020; Holtzman et al., 2005), scRNA-seq detected infection-induced proliferation of ECs and alveolar type 2 (AT2) cells, as well as aberrant appearance of *Trp63*-expressing basal-like cells (Fig. 4A, B), reminiscent of pods or lineage-negative epithelial progenitors observed upon severe H1N1 virus infection (Kumar et al., 2011; Vaughan et al., 2015; Zuo et al., 2015).

**Figure 4:**
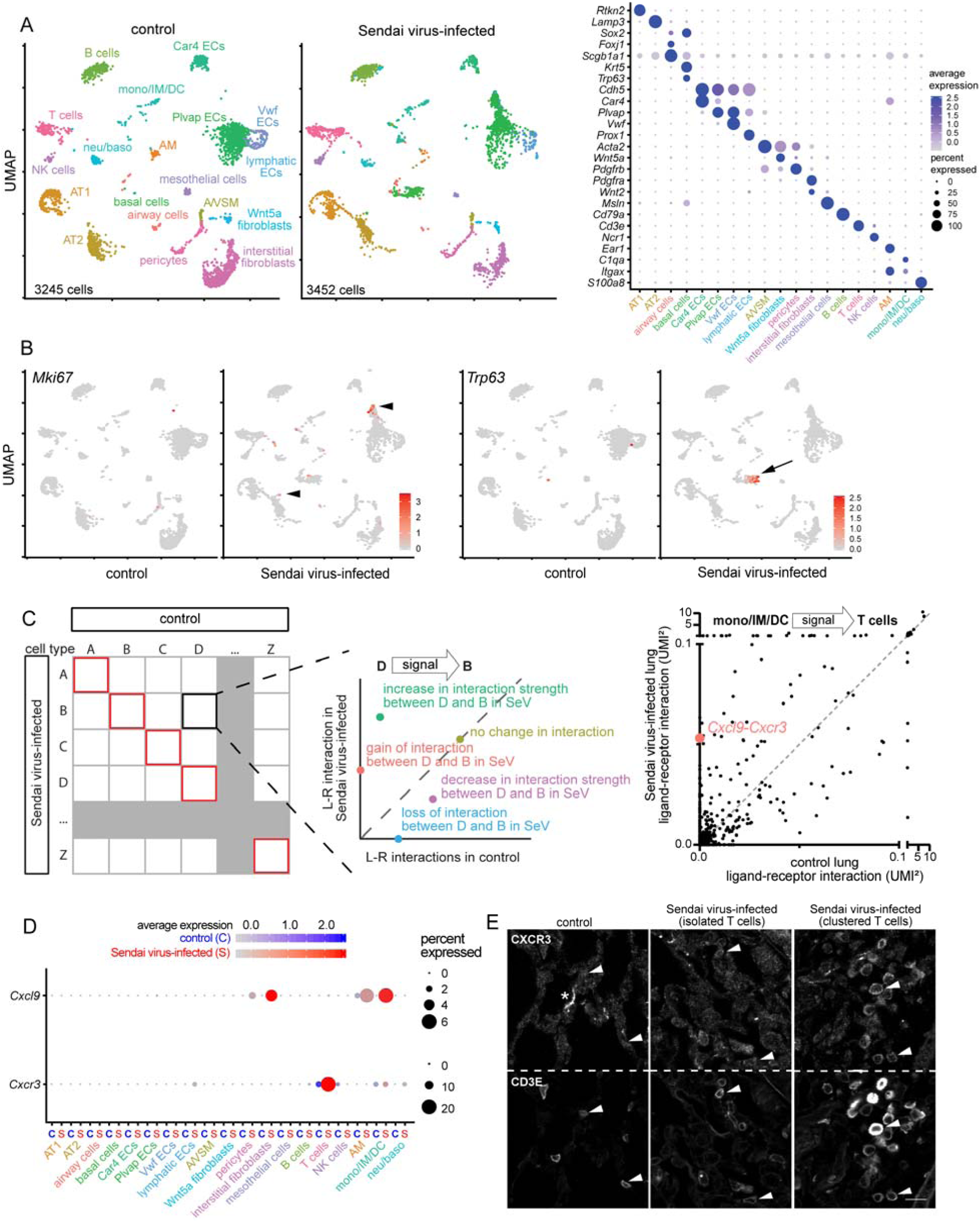
Interactome analysis identifies outlying changes in interaction upon Sendai virus infection. (**A**) Uniform Manifold Approximation and Projections (UMAP) plots (left) of scRNA-seq of control and Sendai virus-infected lungs at 2 weeks post-infection showing 18 cell types, as identified by their marker genes in a dot plot (right). (**B**) Feature plots showing increased proliferation in AT2 cells and ECs (*Mki67*; arrowhead), as well as basal-like cells (*Trp63*; arrow) upon infection. (**C**) Schematic illustrating comparison of control and Sendai virus-infected lungs for each directional cell type-pair. Autocrine interactions are boxed in red along the diagonal. A hypothetical cell type-pair (D to B) illustrates the locations of possible changes in interactions, as demonstrated in a real example showing 2 color-coded interactions. Cell types are abbreviated as in Fig. 2D. (**D**) Dot plots showing upregulation of *Cxcl9* and *Cxcr3* upon infection. (**E**) Confocal images of immunostained control and Sendai virus-infected lungs showing upregulation of CXCR3 in T cells (CD3E), especially when clustered possibly as a result of chemotaxis via CXCR3 (arrows). Asterisk: non-specific extracellular matrix staining. Scale: 10 um.

For each directional cell type pair, we compared control and infected lungs by plotting the corresponding numeric interactions of individual ligand-receptor pairs, such that data points above or below the diagonal line represented enhanced or diminished interactions, respectively, whereas those on the y or x axis represented de novo or lost interactions, respectively. One example was an infection-induced interaction between myeloid cells and T cells via *Cxcl9* and *Cxcr3* (Fig. 4C, D, Table S5). This was supported by CXCR3 immunostaining, showing its upregulation in clustered T cells possibly as a result of chemotaxis (Groom and Luster, 2011; Weng et al., 1998) (Fig. 4E).

To quantify infection-induced changes in interactions, we resorted to the aforementioned concept of outliers by pooling the differences in a given ligand-receptor interaction across 324 cell type pairs and used the same 3 times interquartile range cutoffs (Table S5). We found that 3.6% (27,611 out of 763,344) interactions were outliers, involving 448 and 433 unique ligands and receptors in 289 cell type pairs and averaging 85 outlying interactions per cell type pair (Table S5).

### Integrate interactomes with signaling pathway analysis

To corroborate our interactomes and capitalize on the transcriptomic information available from scRNA-seq, we sought to analyze the activities of signaling pathways and integrate them with associated ligand-receptor pairs. We chose signaling pathways annotated by Gene Ontology, instead of Ingenuity Pathway Analysis, because of its public availability and inclusiveness. Although these databases did not take into account specific biological contexts, limiting their use in general, we reasoned that by averaging over a sufficient number of bona fide, context-independent pathway members, a pathway signature might be still detectible, as assumed in gene set enrichment analysis (Subramanian et al., 2005).

We considered all genes in a pathway as a metagene and generated the corresponding metagene (i.e. pathway) score for each cell, which was then averaged over each cell type (Fig. 5A, Table S6). Using the same outlier concept, albeit with a less stringent cutoff of 1.5 times the interquartile range due to the limited number of cell types available, we pooled individual pathway scores, as well as their changes upon infection, across 18 cell types, and identified outlying pathways (Table S7, S8). We identified 4% (276 out of 6,300) outlying pathways at baseline and 5% (321 out of 6,300) outlying alterations upon infection. To integrate with our interactomes, for a given outlying ligand-receptor pair, we required the receptor to be a member of an outlying signaling pathway in the same cell type as the receiving one in the interactome (Fig. 5A). This integration led to 0.3% (2,323 out of 763,344) outlying interactions at baseline and 0.7% (5,516 out of 763,344) outlying alterations upon infection that were also supported by the corresponding pathway activation (Table S9, S10). One example was infection-enhanced *Bdnf*-*Ntrk2* signaling from AT1 cells to ECs, largely driven by *Ntrk2* upregulation (Fig. 5B). It was tempting to speculate that the supposedly angiogenic role of this signaling (Dalton et al., 2015; Kermani and Hempstead, 2007) contributed to the infection-induced EC proliferation (Fig. 4B).

**Figure 5:**
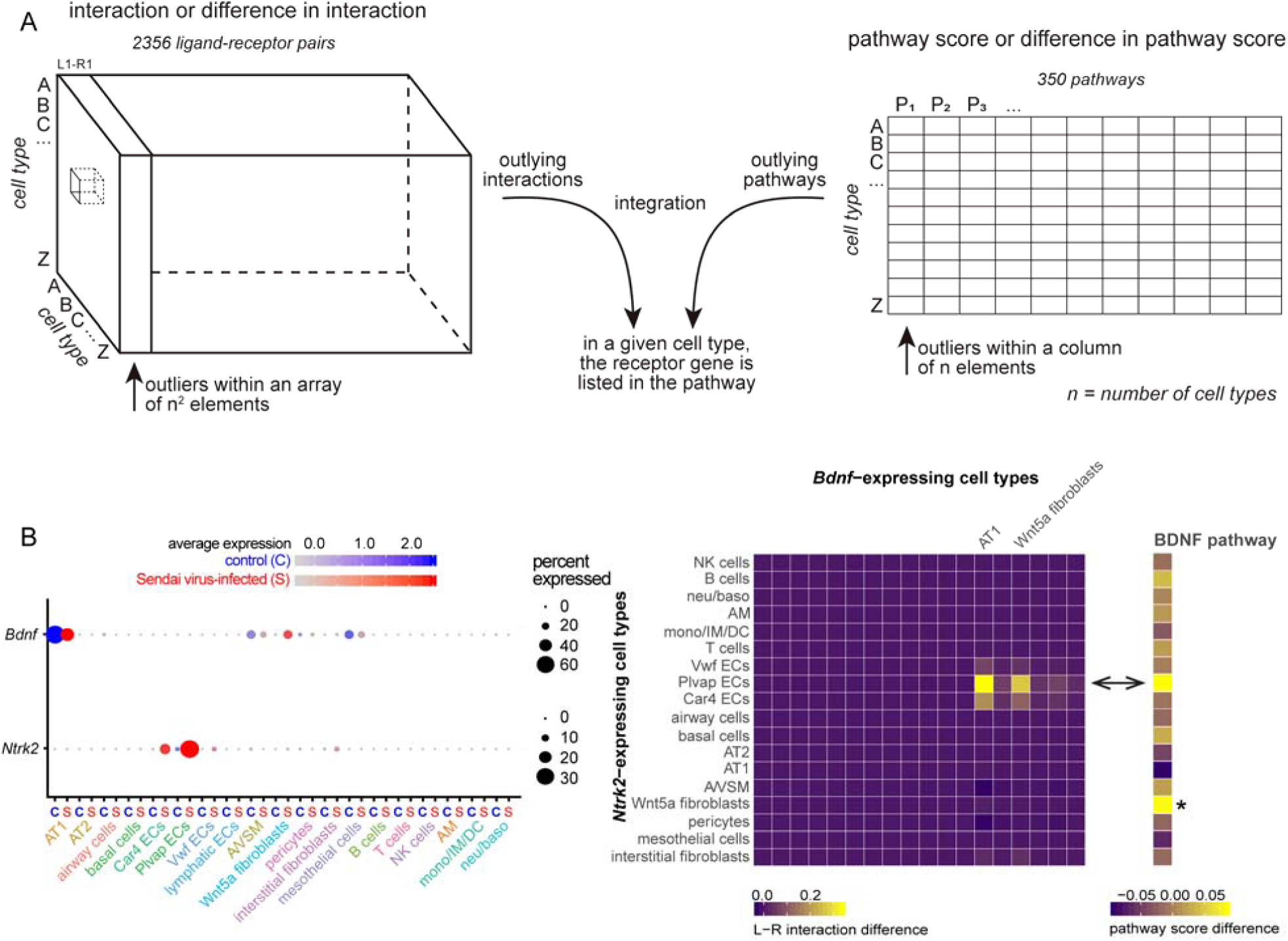
Integration of interactome and pathway scores to prioritize interactions. (**A**) Schematic illustrating the integration of outlying interactions (left) and outlying pathways (right) occurring within the same receiving cell type. Common outliers in interactions/pathways or their changes have the receptor gene included in the pathway gene list in the same receiving cell type. (**B**) Dot plots and heatmaps illustrating a real example of *Bdnf*-*Ntrk2* being both an outlying interaction and an outlying pathway in ECs (double arrowhead). The BDNF pathway is also outlying for the Wnt5a fibroblast population upon infection (asterisk) – likely driven by increased *Bdnf* expression (dot plot), which is not identified as an outlying ligand-receptor interaction as expected from our algorithm, demonstrating the utility of the integrated analysis.

## DISCUSSION

We have optimized experimental and computational methods to construct single-cell interactomes in lungs at baseline and upon infection. Our approaches are intuitive and readily adaptable by bench scientists to study other organs and species, and improve upon existing interactomes (Camp et al., 2017; Cohen et al., 2018; Kumar et al., 2018; Raredon et al., 2019; Skelly et al., 2018; Vento-Tormo et al., 2018; Vieira Braga et al., 2019) in the following aspects.

First, we systematically evaluate and then capitalize on our observations that, on the cell type level, scRNA-seq has comparable sensitivity to genetic driver-based bulk RNA-seq and yet is more robust in identifying and purifying a given cell type (Fig. 1). Our cell type-level analysis bypasses the gene dropout issue inherent to scRNA-seq and imputation methods that are still under development (Gong et al., 2018; Hicks et al., 2018; Huang et al., 2018; Kharchenko et al., 2014; van Dijk et al., 2018). Experimentally, our normalization method is guided by computational assessment of the minimal cell number needed and captures all major lung cell types without inhibitory cost or resorting to non-commercial platforms. A potential concern about cell type-level analysis is failure to capture cellular heterogeneity; this, however, can be alleviated by careful cell type identification. Furthermore, a cell type that is not sufficiently distinct or abundant may not be reliably analyzed even with individual-cell-level algorithms, as evidenced by the significant cell-cell variation of lymphatic ECs (Fig. 1C).

Second, we assign significance based on the concept of outliers, made possible by our numeric representation of ligand-receptor interactions and our normalization protocol to profile a large number of cell types. Alternatively, significance was computed by permutation to evaluate observed versus expected expression in individual cells and thresholded to a binary outcome, which was counted or cross-correlated to represent the interaction strength between cells (Vento-Tormo et al., 2018; Vieira Braga et al., 2019). However, counting interactions may miss functional interactions that are few in number; thresholding at an early step may introduce irremediable bias. Our analysis preserves numeric representations of interactions and pathway scores so that users can define and adjust cutoffs afterwards – 1.5 or 3 times the interquartile range in this case. In addition, using other cell types as internal controls to identify outliers mimics in vivo competition among cell types to achieve specific signaling. Independent from our work, numeric interaction scores can also be used to correlate with other biological scores (Kumar et al., 2018).

Last, we supplement our interactomes with signaling pathway scores to allay the concern of inferring functionality solely based on ligand-receptor expression. Prior knowledge, including the ligand-receptor pair list and pathway components collected by Gene Ontology or Ingenuity Pathway Analysis, is necessary for bioinformatics but largely ignores specific biological contexts; the resulting analysis often errs on the side of false positives. Integrating interactomes with pathway scores, both of which likely include false positives, allows for prioritizing candidate genes. The accuracy may be further improved by better knowledge of ligand-receptor biology. One effort was to consider the multi-component ligand-receptor complex and assume the lowest-expressing component as rate-limiting (Vento-Tormo et al., 2018), which, however, might introduce false negatives. Significant improvement is likely to follow elucidation of signal transduction components and transcriptional targets of individual ligand-receptor pairs.

## METHODS

### Sendai virus infection

Sendai virus (parainfluenza type 1) strain 52 (ATCC #VR-105, RRID:SCR_001672) was expanded in primary rhesus monkey kidney cells (Cell Pro Labs #103-175). Wild-type C57Bl/6 mice obtained from JAX were anaesthetized with isofluorane, suspended by the maxillary incisors on a dosing board with a 60° incline, and infected with a sub-lethal dose of 2.1 × 10^7^ plaque forming units (pfu) of virus in 40 µl phosphate buffered saline (PBS) through oropharyngeal aspiration. All animal experiments were approved by the Institutional Animal Care and Use Committee at Texas A&M Health Science Center Institute of Biosciences and Technology and MD Anderson Cancer Center.

### Section immunostaining and cell counting

For cell lineage counting by immunostaining, lungs were inflated and harvested as previously described (Yang et al., 2016). Briefly, mice were anaesthetized using Avertin (T48402, Sigma) and the lungs perfused with PBS. The trachea was cannulated and the lungs were inflated to full capacity with 0.5% paraformaldehyde (PFA; P6148, Sigma) in PBS at 25 cm H_2_O pressure. The harvested lungs were immersion-fixed in 0.5% PFA at room temperature for 4-6 hr and washed overnight in PBS at 4°C. Section immunostaining was performed following published protocols with minor modifications (Alanis et al., 2014; Chang et al., 2013). Fixed lungs were dissected and cryoprotected overnight at 4°C in 20% sucrose in PBS containing 10% optimal cutting temperature compound (OCT; 4583, Tissue-Tek). The lobes were then frozen in OCT-filled embedding molds on a dry ice and 100% ethanol slurry then kept at −80°C until sectioned. Sections were obtained at 10 um thickness and then blocked in PBS with 0.3% Triton X-100 and 5% normal donkey serum (017-000-121, Jackson ImmunoResearch). Blocked sections were incubated in a humidified chamber at 4 °C overnight with the following primary antibodies diluted in PBS containing 0.3% Triton X-100: GFP (chicken, 1:5,000, Abcam, AB13970), ERG (rabbit, 1:5,000, Abcam, AB92513), CD45 (rat, 1:2,000, eBioscience, 14-0451-81), CD3E (Armenian hamster, 1:250, Biolegend, 100301), and CXCR3 (rat, 1:250, R&D Systems, MAB1685). The sections were submerged in PBS for 30 min and incubated with 4’,6-diamidino-2-phenylindole (DAPI) and donkey secondary antibodies (Jackson ImmunoResearch) diluted in PBS with 0.3% Triton X-100 at room temperature for 1 hr. The sections were washed again in PBS then mounted with Aqua-Poly/Mount (18606, Polysciences) and imaged on a Nikon A1plus confocal microscope or an Olympus FV1000 confocal microscope. For cell lineage counting, 3 images with airway located in the center were taken from 2 *Rosa*^*Sun1GFP/+*^; *Shh*^*Cre/+*^ lungs, in which all epithelial cells were labeled with GFP (Little et al., 2019) to allow costaining for endothelial and immune cells. Endothelial cells (ERG^+^ nuclei) and alveolar epithelial cells (GFP^+^ nuclei) excluding the airways were counted using Fiji’s ‘Find Maxima’ function. Airway epithelial cells were counted by drawing region-of-interest around airways and by using Fiji to Find Maxima for DAPI. Immune cells (CD45^+^ nuclei) and mesenchymal cells (nuclei negative for the other markers) were counted manually.

### Cell dissociation and labeling

Mice were anaesthetized using Avertin injected intraperitoneally, the chest cavity was exposed, and the lungs were perfused with PBS injected through the right ventricle. After clearance of circulating cells, the lungs were removed into PBS and finely minced using forceps. The lung tissue was digested in Leibovitz’s L-15 media (Gibco, 21083-027) with 2 mg/mL Collagenase Type I (Worthington, CLS-1, LS004197), 2 mg/mL Elastase (Worthington, ESL, LS002294), and 0.5 mg/mL DNase I (Worthington, D, LS002007) for 30 min on a 37°C heat block. The digestion was stopped by addition of fetal bovine serum (FBS; Invitrogen, 10082-139) to a final concentration of 20%. The samples were moved to ice and all remaining steps were performed in a cold room. The tissue was triturated at 15 min into digestion and also after digestion was quenched. The resulting cell suspension was filtered through a 70 μm cell strainer (Falcon, 352350), centrifuged at 5,000 rpm for 1 min, and the pellet resuspended in red blood cell lysis buffer (15 mM NH_4_Cl, 12 mM NaHCO_3_, 0.1 mM EDTA, pH 8.0) for 3 min. The cells were pelleted again at 5,000 rpm for 1 min and washed with Leibovitz’s L-15 media with 10% FBS then filtered into a 5 ml tube with a cell strainer cap (Falcon, 352235). The cells were incubated with the following conjugated antibodies: CD45-PE/Cy7 (BioLegend, 103114), ICAM2-A647 (Life tech, A15452), E-cadherin-A488 (Invitrogen, 53-3249-82) at a concentration of 1:250 for 30 min. The cells were centrifuged at 5,000 rpm for 1 min, washed in Leibovitz’s L-15 media with 10% FBS for 5 min and finally resuspended with Leibovitz’s L-15 media with 10% FBS and filtered through a strainer cap. The sample was incubated with SYTOX Blue (Invitrogen, S34857) and sorted on a BD FACSAria Fusion Cell Sorter. The cells were collected in a volume of 250 ul PBS with 10% FBS per collection tube.

### FACS sorting and normalization

After exclusion of dead cells and doublets, cells were gated into the 4 cell lineage populations using a serial gating strategy. All CD45 positive cells were collected as the immune population; CD45 negative cells exhibiting ICAM2 positive signal were collected as endothelial cells; CD45 negative, ICAM2 negative, but E-cadherin positive cells were collected as epithelial cells; cells negative for all markers were collected as mesenchymal cells. To determine cell lineage population-specific reduction in viability after sorting, an equal number of cells from each lineage were mixed and, after adding fresh SYTOX Blue, resorted to determine the percentages of each lineage. These percentages were compared to the expected percentage, 25%, and lineage-specific viability correction factors were calculated and taken into account when combining the 4 lineages for single-cell RNA-seq to achieve as close to equal proportions as possible.

### Single-cell RNA-seq data analysis

#### Data generation

The normalized whole lung samples were processed through the Chromium Single Cell Gene Expression Solution Platform (10x Genomics) using the Chromium Single Cell 3’ Library and Gel Bead Kit (v2, rev D). The library sequencing was performed on an Illumina NextSeq500 using a 26×124 sequencing run format with 8 bp index (Read1). The single-cell reads were aligned against the mm10 mouse reference genome (provided by 10x Genomics), counted, and aggregated using the Cell Ranger pipeline (version 3.0, 10x Genomics). Raw data for the P7 wild-type lung and control and 2 week post-Sendai virus-infected lungs were deposited at GEO under accession numbers GSE144769 and GSE144678.

#### Cell type identification

Data processing was performed using the R package Seurat (version 3.1.2) (Butler et al., 2018). Unless specified, default parameters were used. Single-cell count data used for all downstream calculations was normalized to the number of reads per cell using Seurat’s normalization function (normalized UMI). Data was scaled before undergoing dimensionality reduction using the RunPCA Seurat function. Cells were visualized on a projection map using the RunUMAP function. Feature plots for known lung cell type markers were used to identify the cell types clustered using the Louvain algorithm implemented in the FindClusters function after cells were embedded in a graph using FindNeighbors. The resolution parameter was adjusted to appropriately cluster the cells based on the known cellular gene markers. The cell type distributions were calculated by dividing the number of cells assigned to each cell type by the total number of cells in the sample. The number of unique genes detected in each cell type was counted. Dot plots were used to visualize the marker genes identifying each cell type.

#### Bulk RNA-seq versus scRNA-seq comparison

To calculate the dropout rate of genes in the scRNA-seq data, normalized UMI averaged across all cells for a given cell type was compared to gene expression (FPKM) from the corresponding bulk RNA-seq data (AT1 and AT2 cells from GSE129583; endothelial cells from GSE124324). For genes within specified bins of bulk RNA-seq expression values, the fraction of genes with zero expression in scRNA-seq was calculated and plotted.

#### Analysis of the number of genes expressed per cell type

The number of genes expressed by each cell of one cell type was counted. The cells were ordered both by increasing and by decreasing number of genes expressed. Then, as one cell was added after another, the number of unique genes detected in that cell not detected in the previous cells was tallied, resulting in the cumulative number of unique genes detected for that cell type.

#### Interactome analysis

The normalized ligand and receptor gene count data was accessed from the Seurat object, averaged over all the cells of each cell type, and plotted for each cell type pair. For each ligand-receptor pair for each cell type pair, we multiplied the average ligand and receptor gene expression. The heatmap visualizing these values was created using ggplot2 (version 3.2.1). To identify outlying interactions, we computed the interquartile range (IQR) across all cell type pairs for each ligand-receptor interaction and used a cutoff of 3 times IQR.

#### GO pathway activation score

The 350 GO Pathway gene lists publicly available (Gene Ontology Consortium; containing the phrase “SIGNALING_PATHWAY” in the ‘Biological Process’ category) were used to compute pathway activation scores for each cell type using the AddModuleScore function in Seurat. Outlier pathways were identified for each cell type by applying a 1.5 times IQR cutoff across the scores for all cells for each pathway. To identify possible pathway activation as an outcome of a signaling interaction, the gene list for each outlying pathway was used to match the receptor genes of outlying interactions for the same cell type as the signal receiving cell.

## ACKNOWLEDGEMENT

We thank Odemaris Narvaez del Pilar for generating the scRNA-seq data of the postnatal day 7 mouse lung. The University of Texas MD Anderson Cancer Center DNA Analysis Facility and Flow Cytometry and Cellular Imaging Core Facility are supported by the Cancer Center Support Grant (CA #16672). This work was supported by the University of Texas MD Anderson Cancer Center Start-up and Retention Fund, and National Institutes of Health R01HL130129 (JC).

## AUTHOR CONTRIBUTIONS

MPC and JC designed and performed research, and wrote the paper; BJH performed research; all authors read and approved the paper.

## DECLARATION OF INTERESTS

The authors declare no competing interests.

## SUPPLEMENTAL INFORMATION

**Supplemental Figure 1:**
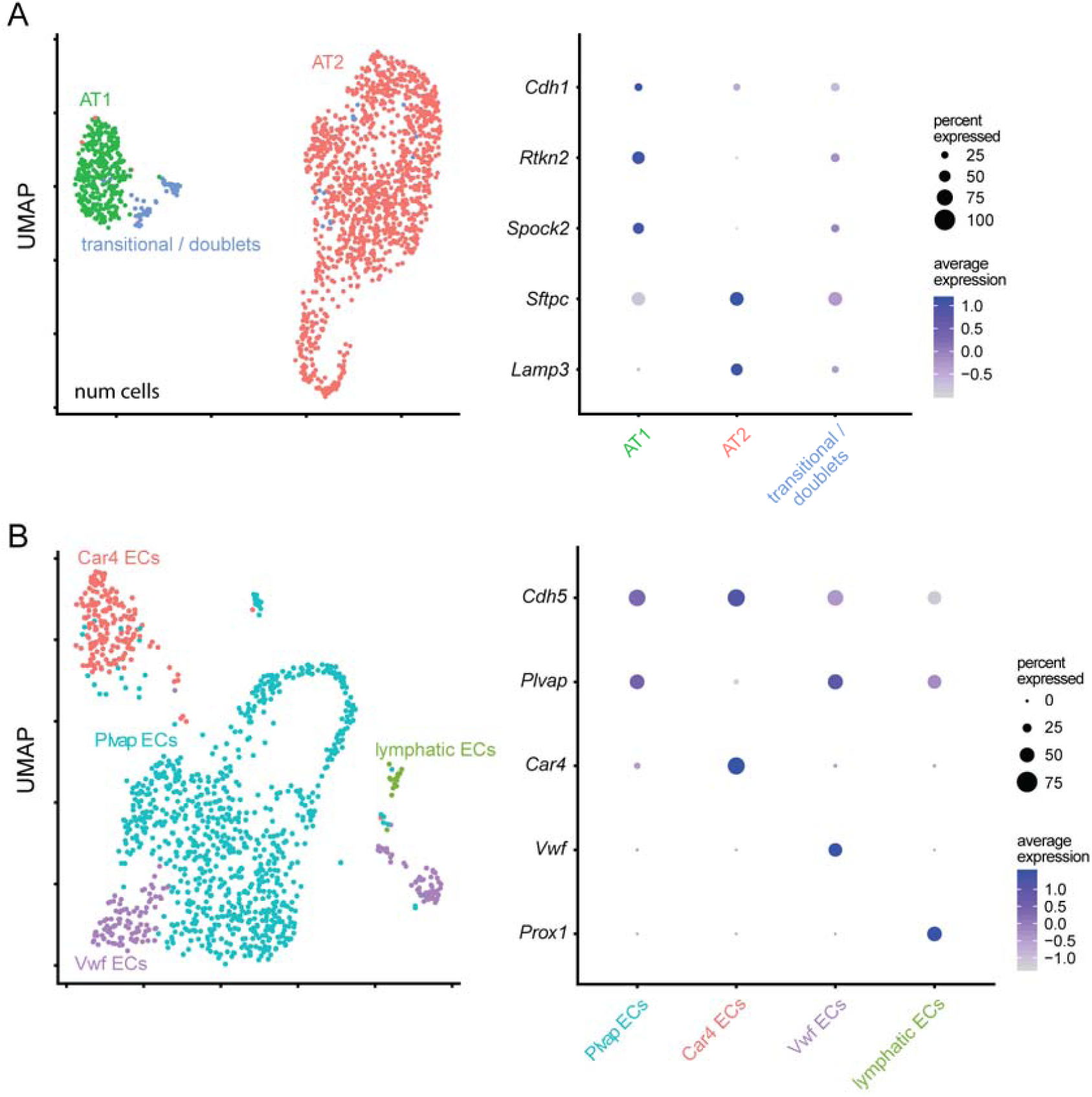
(**A**) Uniform Manifold Approximation and Projections (UMAP) plots (left) of scRNA-seq of P7 lung epithelial cells, as identified by their marker genes in a dot plot (right). Transitional/doublets express both AT1 and AT2 marker genes and are not included in subsequent analysis. (**B**) UMAP plots (left) of scRNA-seq of P7 lung endothelial cell populations, as identified by their marker genes in a dot plot (right).

### Supplemental Tables

**Table S1**. Comparison of bulk RNA-seq and scRNA-seq gene expression in AT1, AT2, and endothelial cell populations.

**Table S2**. Average ligand and receptor gene expression for the 18 cell types of control (worksheet 1) and Sendai virus-infected (worksheet 2) lungs.

**Table S3**. Ligand-receptor interaction scores in each cell type pair in control (worksheet 1) lung and Sendai virus-infected (worksheet 2) lungs.

**Table S4**. Binary indicators of outlying ligand-receptor interactions in each cell type pair in control (worksheet 1) and Sendai virus-infected (worksheet 2) lungs. 1 indicates outlying interaction.

**Table S5**. Differences in ligand-receptor interaction in each cell type pair between control and infected lungs (worksheet 1) and their binary indicators (worksheet 2; 1, increased; −1, decreased; 0, no change).

**Table S6**. GO pathway scores of each cell type in control (worksheet 1) lung and Sendai virus-infected (worksheet 2) lungs.

**Table S7**. Binary GO pathway indicators of each cell type in control (worksheet 1) lung and Sendai virus-infected (worksheet 2) lungs. 1 indicates outlying pathways.

**Table S8**. Differences in GO pathway scores in each cell type between control and infected lungs (worksheet 1) and their binary indicators (worksheet 2; 1, increased; −1, decreased; 0, no change).

**Table S9**. Integration of outlying ligand-receptor interactions and GO pathways in each cell type (individual worksheets) in the control lung.

**Table S10**. Integration of outlying differences in ligand-receptor interactions and GO pathways in each cell type (individual worksheets) between control and Sendai virus-infected lungs.

## REFERENCES

Alanis, D.M., Chang, D.R., Akiyama, H., Krasnow, M.A., and Chen, J. (2014). Two nested developmental waves demarcate a compartment boundary in the mouse lung. Nat Commun 5, 3923.

Angelidis, I., Simon, L.M., Fernandez, I.E., Strunz, M., Mayr, C.H., Greiffo, F.R., Tsitsiridis, G., Ansari, M., Graf, E., Strom, T.M., et al. (2019). An atlas of the aging lung mapped by single cell transcriptomics and deep tissue proteomics. Nat Commun 10, 963.

Butler, A., Hoffman, P., Smibert, P., Papalexi, E., and Satija, R. (2018). Integrating single-cell transcriptomic data across different conditions, technologies, and species. Nat Biotechnol 36, 411–420.

Camp, J.G., Sekine, K., Gerber, T., Loeffler-Wirth, H., Binder, H., Gac, M., Kanton, S., Kageyama, J., Damm, G., Seehofer, D., et al. (2017). Multilineage communication regulates human liver bud development from pluripotency. Nature 546, 533–538.

Chang-Panesso, M., and Humphreys, B.D. (2017). Cellular plasticity in kidney injury and repair. Nat Rev Nephrol 13, 39–46.

Chang, D.R., Martinez Alanis, D., Miller, R.K., Ji, H., Akiyama, H., McCrea, P.D., and Chen, J. (2013). Lung epithelial branching program antagonizes alveolar differentiation. Proc Natl Acad Sci U S A 110, 18042–18051.

Cohen, M., Giladi, A., Gorki, A.D., Solodkin, D.G., Zada, M., Hladik, A., Miklosi, A., Salame, T.M., Halpern, K.B., David, E., et al. (2018). Lung Single-Cell Signaling Interaction Map Reveals Basophil Role in Macrophage Imprinting. Cell 175, 1031–1044 e1018.

Dalton, J.E., Glover, A.C., Hoodless, L., Lim, E.K., Beattie, L., Kirby, A., and Kaye, P.M. (2015). The neurotrophic receptor Ntrk2 directs lymphoid tissue neovascularization during Leishmania donovani infection. PLoS Pathog 11, e1004681.

Goldblatt, D.L., Flores, J.R., Valverde Ha, G., Jaramillo, A.M., Tkachman, S., Kirkpatrick, C.T., Wali, S., Hernandez, B., Ost, D.E., Scott, B.L., et al. (2020). Inducible epithelial resistance against acute Sendai virus infection prevents chronic asthma-like lung disease in mice. Br J Pharmacol.

Gong, W., Kwak, I.Y., Pota, P., Koyano-Nakagawa, N., and Garry, D.J. (2018). DrImpute: imputing dropout events in single cell RNA sequencing data. BMC Bioinformatics 19, 220.

Groom, J.R., and Luster, A.D. (2011). CXCR3 in T cell function. Exp Cell Res 317, 620–631.

Han, X., Wang, R., Zhou, Y., Fei, L., Sun, H., Lai, S., Saadatpour, A., Zhou, Z., Chen, H., Ye, F., et al. (2018). Mapping the Mouse Cell Atlas by Microwell-Seq. Cell 172, 1091–1107 e1017.

Hicks, S.C., Townes, F.W., Teng, M., and Irizarry, R.A. (2018). Missing data and technical variability in single-cell RNA-sequencing experiments. Biostatistics 19, 562–578.

Holtzman, M.J., Tyner, J.W., Kim, E.Y., Lo, M.S., Patel, A.C., Shornick, L.P., Agapov, E., and Zhang, Y. (2005). Acute and chronic airway responses to viral infection: implications for asthma and chronic obstructive pulmonary disease. Proc Am Thorac Soc 2, 132–140.

Huang, M., Wang, J., Torre, E., Dueck, H., Shaffer, S., Bonasio, R., Murray, J.I., Raj, A., Li, M., and Zhang, N.R. (2018). SAVER: gene expression recovery for single-cell RNA sequencing. Nat Methods 15, 539–542.

Jurisic, G., Iolyeva, M., Proulx, S.T., Halin, C., and Detmar, M. (2010). Thymus cell antigen 1 (Thy1, CD90) is expressed by lymphatic vessels and mediates cell adhesion to lymphatic endothelium. Exp Cell Res 316, 2982–2992.

Kermani, P., and Hempstead, B. (2007). Brain-derived neurotrophic factor: a newly described mediator of angiogenesis. Trends Cardiovasc Med 17, 140–143.

Kharchenko, P.V., Silberstein, L., and Scadden, D.T. (2014). Bayesian approach to single-cell differential expression analysis. Nat Methods 11, 740–742.

Kumar, M.P., Du, J., Lagoudas, G., Jiao, Y., Sawyer, A., Drummond, D.C., Lauffenburger, D.A., and Raue, A. (2018). Analysis of Single-Cell RNA-Seq Identifies Cell-Cell Communication Associated with Tumor Characteristics. Cell Rep 25, 1458–1468 e1454.

Kumar, P.A., Hu, Y., Yamamoto, Y., Hoe, N.B., Wei, T.S., Mu, D., Sun, Y., Joo, L.S., Dagher, R., Zielonka, E.M., et al. (2011). Distal airway stem cells yield alveoli in vitro and during lung regeneration following H1N1 influenza infection. Cell 147, 525–538.

Little, D.R., Gerner-Mauro, K.N., Flodby, P., Crandall, E.D., Borok, Z., Akiyama, H., Kimura, S., Ostrin, E.J., and Chen, J. (2019). Transcriptional control of lung alveolar type 1 cell development and maintenance by NK homeobox 2-1. Proc Natl Acad Sci U S A 116, 20545–20555.

Mo, A., Mukamel, E.A., Davis, F.P., Luo, C., Henry, G.L., Picard, S., Urich, M.A., Nery, J.R., Sejnowski, T.J., Lister, R., et al. (2015). Epigenomic Signatures of Neuronal Diversity in the Mammalian Brain. Neuron 86, 1369–1384.

Raredon, M.S.B., Adams, T.S., Suhail, Y., Schupp, J.C., Poli, S., Neumark, N., Leiby, K.L., Greaney, A.M., Yuan, Y., Horien, C., et al. (2019). Single-cell connectomic analysis of adult mammalian lungs. Sci Adv 5, eaaw3851.

Reyfman, P.A., Walter, J.M., Joshi, N., Anekalla, K.R., McQuattie-Pimentel, A.C., Chiu, S., Fernandez, R., Akbarpour, M., Chen, C.I., Ren, Z., et al. (2019). Single-Cell Transcriptomic Analysis of Human Lung Provides Insights into the Pathobiology of Pulmonary Fibrosis. Am J Respir Crit Care Med 199, 1517–1536.

Skelly, D.A., Squiers, G.T., McLellan, M.A., Bolisetty, M.T., Robson, P., Rosenthal, N.A., and Pinto, A.R. (2018). Single-Cell Transcriptional Profiling Reveals Cellular Diversity and Intercommunication in the Mouse Heart. Cell Rep 22, 600–610.

Strunz, M., Simon, L.M., Ansari, M., Mattner, L.F., Angelidis, I., Mayr, C.H., Kathiriya, J., Yee, M., Ogar, P., Sengupta, A., et al. (2019). bioRxiv 705244 [Preprint]. 17 July 2019. https://doi.org/10.1101/705244

Subramanian, A., Tamayo, P., Mootha, V.K., Mukherjee, S., Ebert, B.L., Gillette, M.A., Paulovich, A., Pomeroy, S.L., Golub, T.R., Lander, E.S., et al. (2005). Gene set enrichment analysis: a knowledge-based approach for interpreting genome-wide expression profiles. Proc Natl Acad Sci U S A 102, 15545–15550.

Tabula Muris, C., Overall, c., Logistical, c., Organ, c., processing, Library, p., sequencing, Computational data, a., Cell type, a., Writing, g., et al. (2018). Single-cell transcriptomics of 20 mouse organs creates a Tabula Muris. Nature 562, 367–372.

Tata, P.R., and Rajagopal, J. (2017). Plasticity in the lung: making and breaking cell identity. Development 144, 755–766.

van Dijk, D., Sharma, R., Nainys, J., Yim, K., Kathail, P., Carr, A.J., Burdziak, C., Moon, K.R., Chaffer, C.L., Pattabiraman, D., et al. (2018). Recovering Gene Interactions from Single-Cell Data Using Data Diffusion. Cell 174, 716–729 e727.

Varga, J., and Greten, F.R. (2017). Cell plasticity in epithelial homeostasis and tumorigenesis. Nat Cell Biol 19, 1133–1141.

Vaughan, A.E., Brumwell, A.N., Xi, Y., Gotts, J.E., Brownfield, D.G., Treutlein, B., Tan, K., Tan, V., Liu, F.C., Looney, M.R., et al. (2015). Lineage-negative progenitors mobilize to regenerate lung epithelium after major injury. Nature 517, 621–625.

Vento-Tormo, R., Efremova, M., Botting, R.A., Turco, M.Y., Vento-Tormo, M., Meyer, K.B., Park, J.E., Stephenson, E., Polanski, K., Goncalves, A., et al. (2018). Single-cell reconstruction of the early maternal-fetal interface in humans. Nature 563, 347–353.

Vieira Braga, F.A., Kar, G., Berg, M., Carpaij, O.A., Polanski, K., Simon, L.M., Brouwer, S., Gomes, T., Hesse, L., Jiang, J., et al. (2019). A cellular census of human lungs identifies novel cell states in health and in asthma. Nat Med 25, 1153–1163.

Vila Ellis, L., Cain, M.P., Hutchison, V., Flodby, P., Crandall, E.D., Borok, Z., Zhou, B., Ostrin, E.J., Wythe, J.D., and Chen, J. (2019). bioRxiv 840033 [Preprint]. 13 November 2019. https://doi.org/10.1101/840033

Weng, Y., Siciliano, S.J., Waldburger, K.E., Sirotina-Meisher, A., Staruch, M.J., Daugherty, B.L., Gould, S.L., Springer, M.S., and DeMartino, J.A. (1998). Binding and functional properties of recombinant and endogenous CXCR3 chemokine receptors. J Biol Chem 273, 18288–18291.

Yang, J., Hernandez, B.J., Martinez Alanis, D., Narvaez del Pilar, O., Vila-Ellis, L., Akiyama, H., Evans, S.E., Ostrin, E.J., and Chen, J. (2016). The development and plasticity of alveolar type 1 cells. Development 143, 54–65.

Yuan, T., Volckaert, T., Chanda, D., Thannickal, V.J., and De Langhe, S.P. (2018). Fgf10 Signaling in Lung Development, Homeostasis, Disease, and Repair After Injury. Front Genet 9, 418.

Zuo, W., Zhang, T., Wu, D.Z., Guan, S.P., Liew, A.A., Yamamoto, Y., Wang, X., Lim, S.J., Vincent, M., Lessard, M., et al. (2015). p63(+)Krt5(+) distal airway stem cells are essential for lung regeneration. Nature 517, 616–620.

